# The impact of rare variation on gene expression across tissues

**DOI:** 10.1101/074443

**Authors:** Xin Li, Yungil Kim, Emily K. Tsang, Joe R. Davis, Farhan N. Damani, Colby Chiang, Zachary Zappala, Benjamin J. Strober, Alexandra J. Scott, Andrea Ganna, Jason Merker, GTEx Consortium, Ira M. Hall, Alexis Battle, Stephen B. Montgomery

**Author notes:** equal contribution (Xin Li, Yungil Kim, Emily K. Tsang, Joe R. Davis).

## Abstract

Rare genetic variants are abundant in humans yet their functional effects are often unknown and challenging to predict. The Genotype-Tissue Expression (GTEx) project provides a unique opportunity to identify the functional impact of rare variants through combined analyses of whole genomes and multi-tissue RNA-sequencing data. Here, we identify gene expression outliers, or individuals with extreme expression levels, across 44 human tissues, and characterize the contribution of rare variation to these large changes in expression. We find 58% of underexpression and 28% of overexpression outliers have underlying rare variants compared with 9% of non-outliers. Large expression effects are enriched for proximal loss-of-function, splicing, and structural variants, particularly variants near the TSS and at evolutionarily conserved sites. Known disease genes have expression outliers, underscoring that rare variants can contribute to genetic disease risk. To prioritize functional rare regulatory variants, we develop RIVER, a Bayesian approach that integrates RNA and whole genome sequencing data from the same individual. RIVER predicts functional variants significantly better than models using genomic annotations alone, and is an extensible tool for personal genome interpretation. Overall, we demonstrate that rare variants contribute to large gene expression changes across tissues with potential health consequences, and provide an integrative method for interpreting rare variants in individual genomes.

## Introduction

The recent and rapid expansion of human populations has led to an abundance of rare genetic variants some of which are expected to contribute to an individual’s genetic risk of disease^1^–^4^. However, prioritizing the subset of rare variants most likely to cause cellular and phenotypic changes from the tens of thousands of rare variants within each individual’s genome remains a major challenge. While genetic association analyses have successfully identified many common genetic risk factors for non-Mendelian traits, rare variants are private or at such low frequency that association studies become infeasible^1^,^5^. To overcome this challenge, multiple approaches for distinguishing pathogenic from benign rare variants have leveraged the genetic code to identify nonsense or other deleterious protein coding alleles^1^,^6^–^8^. Such variants not only inform individual genetic risk but are valuable natural gene knockouts that underlie extreme phenotypes and help predict potential drug targets. Unfortunately, no analogous code exists for identifying non-coding variants with functional consequences.

Promising models have been developed to predict variant impact from diverse genomic features, including cis-regulatory element annotation and conservation status^9^–^13^. We hypothesized that incorporating each individual’s gene expression data would improve prioritization of functional rare variants. Indeed, for rare loss-of-function variants in protein-coding regions, allele-specific effects across multiple tissues have characterized the systemic impact of nonsense-mediated decay^14^,^15^. In single-tissue studies, rare non-coding variants, in aggregate, have been associated with outlier gene expression levels, suggesting their potential to drastically alter gene expression^16^–^19^. However, it remains unknown which categories of rare variation have the strongest impact on gene expression and how their consequences are reflected across multiple tissues. As whole genome sequencing becomes more prevalent, new means to understand rare variant biology and to prioritize the variants with important individual consequences will be essential to personal genomics and its integration in precision medicine.

### Extreme expression is shared across tissues

To assess the impact of rare genetic variation on gene expression in diverse human tissues, we analyzed data from the Genotype Tissue Expression project (GTEx V6p), which includes 7,051 RNA-sequencing samples from 44 tissues in 449 individuals (median of 126 individuals per tissue and 16 tissues sampled per individual). We restricted rare variant analysis to the 123 individuals of European ancestry, but used the entire cohort for all other analyses (Extended Data Fig. 1). We defined rare variants as those with minor allele frequency below 1% within GTEx as well as within the European panel of the 1000 Genomes project for single nucleotide variants (SNVs) and short insertions and deletions (indels)^20^. Each individual had a median of 43,739 rare SNVs, 4,835 rare indels and 59 rare structural variants (SVs) (Extended Data Fig. 2).

Our analysis focused on individuals with extremely high or low expression of a particular gene compared with the rest of the cohort. We refer to these individuals as *gene expression outliers*. The GTEx data affords the ability to identify both *single-tissue* and *multi-tissue expression outliers*, with the latter showing consistent extreme expression for a gene across many tissues. To account for broad environmental or technical confounders, we removed hidden factors estimated by PEER^21^ from each tissue, which increased the predictive power of outlier expression across tissues (Extended Data Fig. 3). After confounder removal and data normalization, we identified both single-tissue and multi-tissue outliers among the entire cohort of 449 individuals. For each tissue, an individual was called a *single-tissue outlier* for a particular gene if that individual had the largest absolute Z-score and the absolute value was at least two. For each gene, the individual with the most extreme median Z-score taken across tissues was identified as a *multi-tissue outlier* for that gene provided the absolute median Z-score was at least two (Fig. 1a). Therefore, each gene had at most one single-tissue outlier per tissue and one multi-tissue outlier. Under this definition an individual can be an outlier for multiple genes.

**Figure 1.**
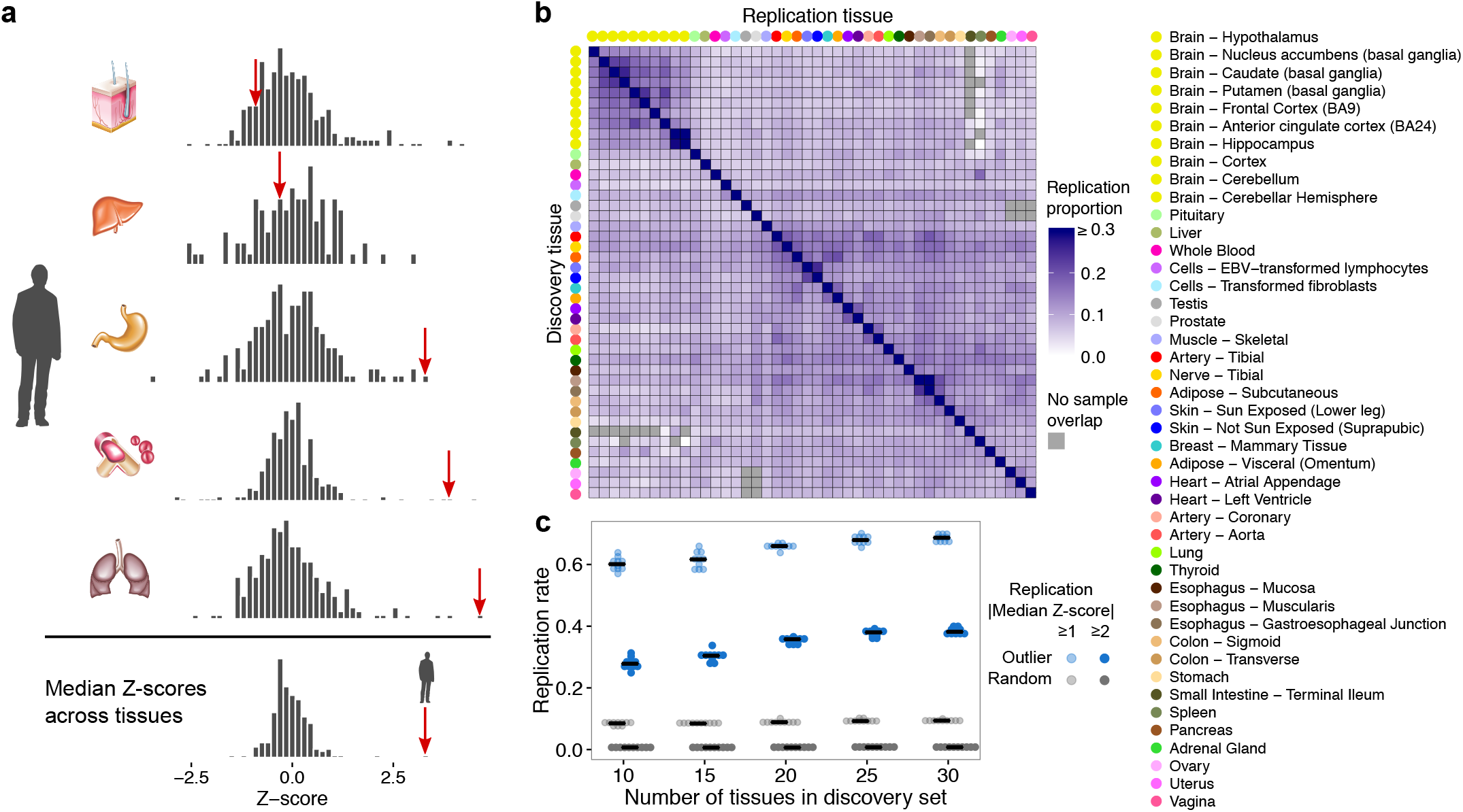
Gene expression outliers and sharing between tissues. (a) A multi-tissue outlier. In this example, the individual has extreme expression values for the gene *AKR1C4* in multiple tissues (red arrows) and the most extreme median expression value across tissues. (b) Outlier expression sharing between tissues, as measured by the proportion of single-tissue outliers that have |Z-score| ≥ 2 for the corresponding genes in each replication tissue. Tissues are hierarchically clustered by gene expression. (c) Estimated replication rate of multi-tissue outliers in a constant held-out set of tissues for different sets of discovery tissues. We compared outliers identified in the discovery set to the same number of randomly selected individuals (see Online methods). Due to incomplete tissue sampling, the number of tissues supporting each outlier is at least five but less than the size of the discovery set.

We identified a single-tissue expression outlier for almost all expressed genes (≥ 99%) in each tissue and a multi-tissue outlier for 4,919 of 18,380 tested genes (27%). Each individual was a single-tissue outlier for a median of 1,653 genes (83 per tissue) compared with a median of 10 genes as a multi-tissue outlier. We confirmed that known environmental factors of race, sex, and BMI were uncorrelated with the number of genes for which an individual was a multi-tissue outlier (Extended Data Fig. 4). We did observe a weak but statistically significant, positive correlation with ischemic time (Spearman *ρ* = 0.175, nominal *P* = 0.00022) and age (Spearman *ρ* = 0.101, nominal *P* = 0.033). Single-tissue outliers discovered in one tissue replicated in other tissues at rates up to 33%, with stronger replication rates among related tissues, such as the two skin tissues as well as the left ventricle and atrial appendage of the heart (Fig. 1b). Replication estimates were underestimated for tissues with smaller sample sizes but biased upward for pairs of tissues with many overlapping individuals sampled (Extended Data Fig. 5). However, we confirmed that the overall sharing patterns were maintained when we accounted for sampling differences, using pairs of tissues with enough overlapping samples to assess the inflation directly. Single-tissue outliers were also detected as multi-tissue outliers at rates from 1.2% to 5.6%, with more overlap for tissues with more samples (Extended Data Fig. 6, Pearson r = 0.79, *P* = 1.4 × 10^−10^). While tissue-specific expression may partially explain the small overlap, the trend is most likely due to the inherent noise in the single-tissue analyses. Indeed, the replication rate for multi-tissue outliers was much higher than for single-tissue outliers and increased with the number of tissues used for discovery, highlighting the value of multiple tissue data for robust outlier detection (Fig. 1c). The difference in replication rate between outliers and randomly selected individuals was greater than could be explained by the bias from overlapping individuals in the discovery and replication sets.

### Functional rare variants underlie multi-tissue outliers

We investigated the extent to which extreme expression could be explained by genetic variation. Here, we focused on the 123 individuals of European descent with whole genome sequencing (average coverage 30X), among whom we identified 1,144 multi-tissue outliers. We evaluated the proportion of outliers with variants at different frequencies within 10 kb of the transcription start site (TSS) compared to corresponding genes in non-outliers to identify the effects of variants acting in cis. Multi-tissue outliers were more enriched for rare variants than common ones (Fig. 2a). This enrichment was most pronounced for structural variants (SVs), and larger for short insertions and deletions (indels) than for single nucleotide variants (SNVs). The enrichment for rare variants was markedly stronger for multi-tissue outliers compared to single-tissue outliers (Fig. 2b, Extended Data Fig. 7), a trend that became more striking at larger Z-score thresholds (Fig. 2b).

**Figure 2.**
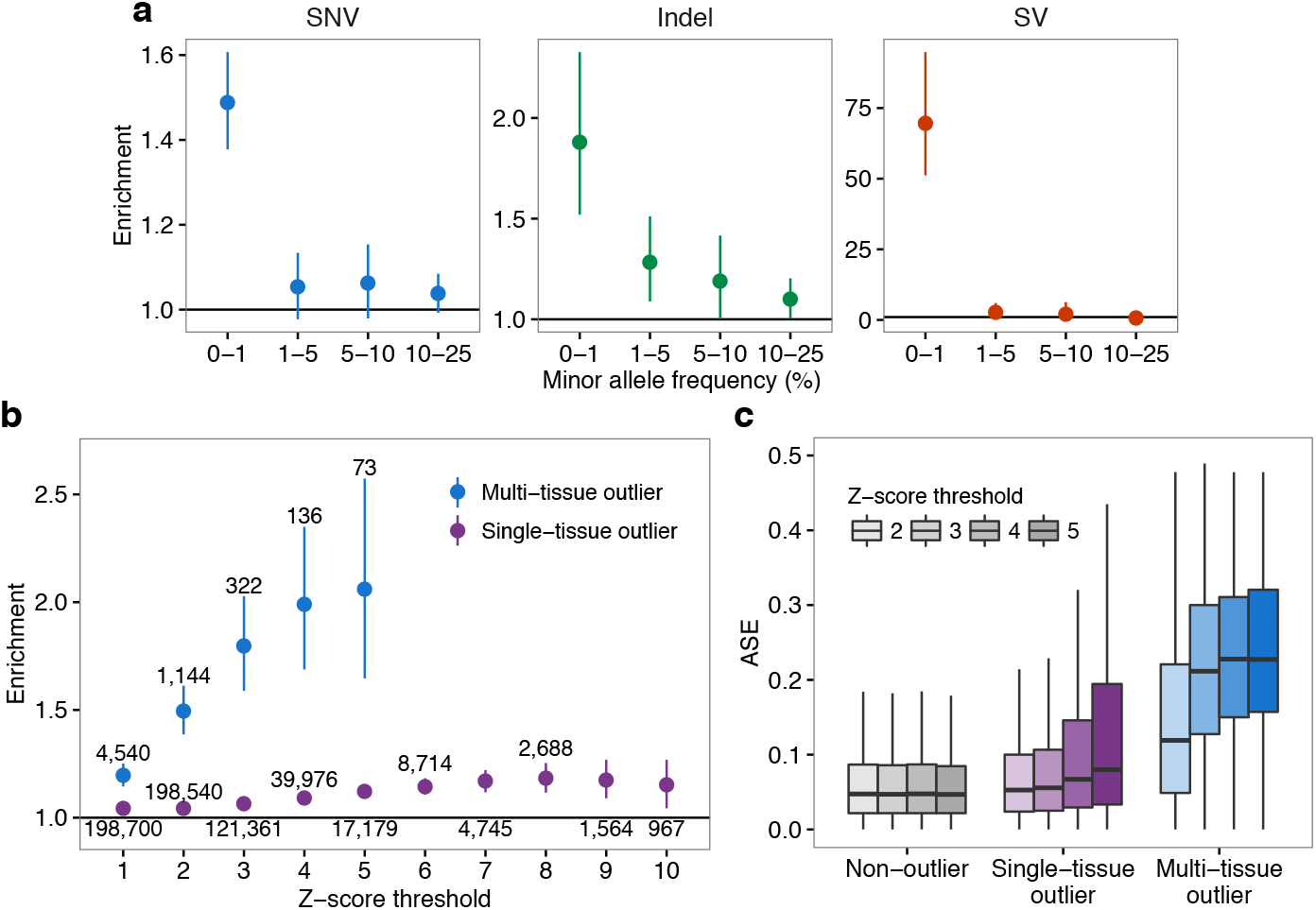
Enrichment of rare variants and ASE in outliers. (a) Enrichment of SNVs, indels, and SVs within 10 kb of the TSS of genes with outliers in the corresponding outlier individuals, as compared with the same genes in non-outlier individuals. For each frequency stratum, we calculated enrichment as a ratio of proportions. The numerator is the proportion of outliers with a variant whose frequency lies within the range, and the denominator is the corresponding proportion for non-outliers. Bars indicate 95% Wald confidence intervals. (b) Rare SNV enrichment for multi-tissue and single-tissue outliers at increasing Z-score thresholds. This threshold applies to the median absolute Z-score for multi-tissue outliers and the absolute Z-score for single-tissue ones. Text labels indicate the number of outliers at each threshold. (c) ASE, measured as the magnitude of the difference between the reference-allele ratio and the null expectation of 0.5. The non-outlier category is defined in the Online methods section.

As rare variants are often heterozygous, expression outliers driven by rare variants in cis should exhibit allele-specific expression (ASE). At multiple Z-score thresholds, both single-tissue and multi-tissue outliers were significantly enriched for ASE, as compared to non-outliers (two-sided Wilcoxon rank sum tests, each nominal *P* < 2.2 × 10^−16^). ASE was stronger for multi-tissue outliers than for single-tissue outliers, and increased with the Z-score threshold (Fig. 2c). This, along with the stronger rare variant enrichments for multi-tissue outliers, suggests that single-tissue outliers are less robust to non-genetic confounders.

We aimed to identify the specific properties of rare genetic variants that induce large changes in gene expression. We evaluated the enrichment of diverse variant classes (Extended Data Table 1) in outliers compared with non-outliers. To capture both coding and non-coding variant classes, we evaluated variants in the gene body and up to 10 kb (200 kb for SVs and variants in enhancers) from the transcription start or end sites of genes with outliers. SVs, taken together, had the strongest enrichment, and their impact on gene expression across tissues is well characterized^22^. We also observed, in order of significance, enrichments for variants near splice sites, introducing frameshifts, at start or stop codons, near the TSS, outside of coding regions and among the top 1% of CADD or vertebrate PhyloP scores, and with other coding annotations (Fig. 3a). These results suggest that variants in coding regions contribute disproportionately to outlier expression. Indeed, we observed weakened enrichments for all variants types (SNVs, indels, and SVs) when excluding exonic regions (Extended Data Fig. 8).

**Figure 3.**
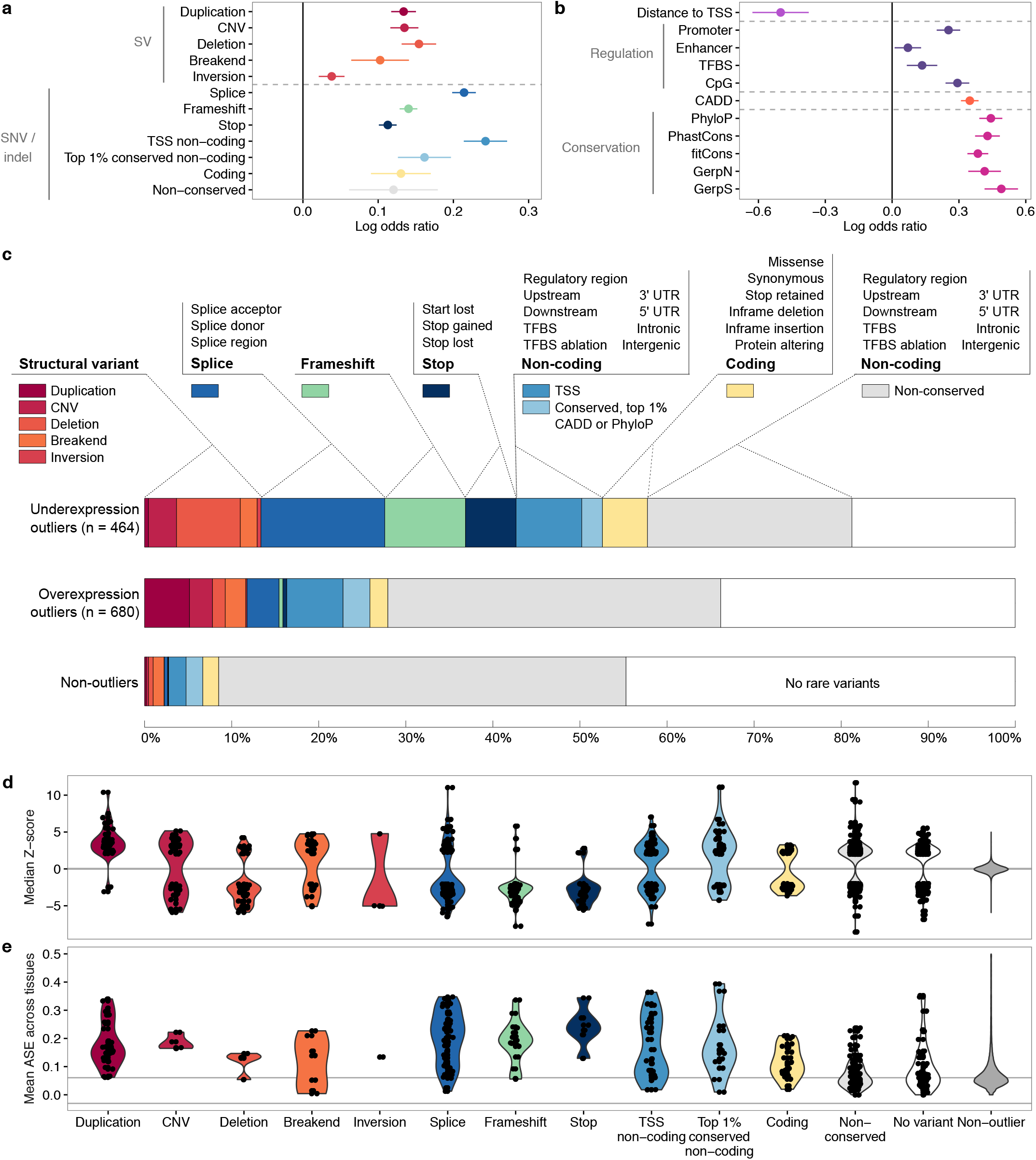
Stratification of multi-tissue outliers by rare variant classes. (a) Enrichment of disjoint variant classes among outliers. Log odds ratio and 95% Wald CI calculated for each variant class by fitting a univariate logistic regression model of outlier status versus the variant class. We defined regions near transcription start sites (TSS) as 250 bp upstream to 750 bp downstream of the most upstream annotated TSS. For each gene, we considered rare variants in the gene body and within 10 kb of either end of the gene (200 kb for SVs). (b) Enrichment of functional annotations for rare SNVs and indels within 10 kb of the gene, including the gene body (200 kb for enhancers). (c) Proportion of genes with an outlier potentially explained by each rare variant class. Each outlier gene is counted, at most once, for the variant class with the most significant enrichment (as ordered from left to right). (d) Distribution of median Z-scores for each variant class. Positive values represent overexpression and negative values underexpression. (e) For each variant class, distribution of ASE effect sizes, the absolute difference between the reference-allele ratio and 0.5, averaged across tissues. Grey lines mark the median values among non-outliers.

To identify the relationship between outlier expression and genomic annotation, we tested whether rare variants near genes with outliers had high conservation or CADD scores^9^ and whether they occurred in known regulatory regions. Multi-tissue outliers were strongly enriched for variants in promoter or CpG sites, and they had variants with higher conservation and CADD scores than non-outliers. We observed a weaker enrichment for variants in enhancers and transcription factor binding sites (Fig. 3b, Extended Data Fig. 9). By jointly considering major classes of variation, we observed that 58% of underexpression and 28% of overexpression outliers had rare variants near the relevant gene, compared with 9% for non-outliers (Fig. 3c). These results confirmed that rare variation is more likely to decrease expression^23^–^25^ and that overexpression outliers may more often be due to environmental factors. Some variant classes had strong directionality in their effect: duplications caused overexpression outliers, while deletions, start and stop codon variants, and frameshifts led to underexpression outliers (Fig. 3d). This directionality agrees with the expected regulatory effect of these variant types and offers further evidence for the role of genetic variation in outlier expression. There was also strong ASE for outliers carrying all categories of variants except those with only non-conserved variants or without any rare variants near the gene (Fig. 3e), which suggests that common variants or non-genetic factors likely caused the extreme expression in those cases.

### Constrained genes rarely have multi-tissue outliers

We hypothesized that rare functional variants and extreme expression in essential genes would be subject to selective pressure. Consistent with ongoing purifying selection against large, multi-tissue effects, rare promoter variants in outliers exhibited significantly lower allele frequencies in the UK10K cohort of 3,781 individuals^3^ than those in non-outliers for the same genes (Fig. 4a, two-sided Wilcoxon rank sum test, *P* = 0.0013). Genes intolerant to loss-of-function mutations as curated by the Exome Aggregation Consortium^26^ were depleted of multi-tissue outliers and multi-tissue eQTLs (Fisher’s exact test, both *P* < 2.2 × 10^−16^; Fig. 4b), which supports our hypothesis that altering expression levels of critical genes can be deleterious. We observed a similar depletion in genes resistant to missense variation (for genes with outliers *P* = 1.676 × 10^−15^ and for multi-tissue eGenes *P* < 2.2 × 10^−16^; Extended Data Fig. 10a). Genes with a multi-tissue outlier were enriched for multi-tissue eQTLs (two-sided Wilcoxon rank sum test *P* < 2.2 × 10^−16^, Extended Data Fig. 10c,d). However, we found some evidence that genes with outliers were more constrained for missense and loss-of-function variation than genes with multi-tissue eQTLs (Tukey’s range test, missense Z-score *P* = 0.0044, probability of loss-of-function intolerance score *P* = 0.086; Fig. 4b, Extended Data Fig. 10a).

**Figure 4.**
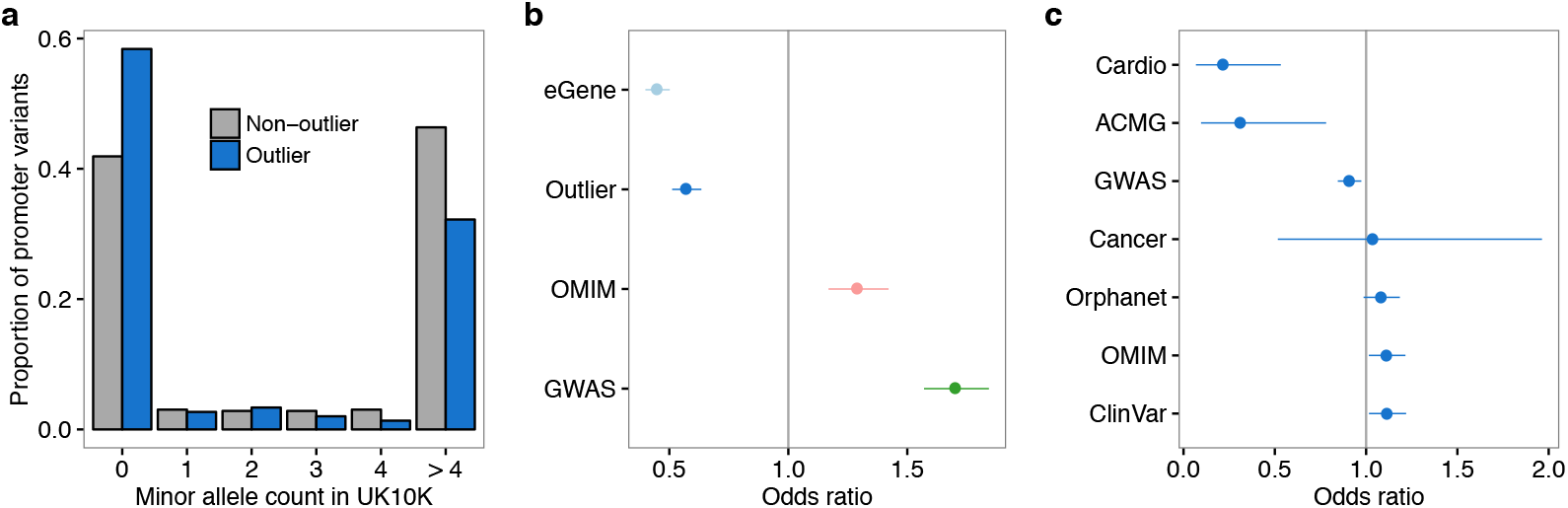
Evolutionary constraint of genes with multi-tissue outliers. (a) Distributions of UK10K minor allele frequencies for promoter SNVs in outlier and non-outlier individuals at genes with multi-tissue outliers. (b) Odds ratio of being intolerant to loss-of-function variants, as defined by ExAC, for genes with multi-tissue outliers, genes with shared eQTLs (eGene), genes reported in the GWAS catalog, and OMIM genes. Bars represent 95% confidence intervals (Fisher’s exact test). (c) Depletion of disease genes among genes with multi-tissue outliers. Odds ratio of a gene having a multi-tissue outlier for each of eight sets of genes involved in complex traits or diseases. Bars represent 95% confidence intervals (Fisher’s exact test).

We expected disease genes to be depleted of multi-tissue expression outliers in the general population since extreme expression at critical genes may have severe health consequences. We confirmed this among the GTEx individuals for two well curated disease gene lists: a list of genes involved in heritable cardiovascular disease (Cardio) and genes in the ACMG guidelines for incidental findings (Fig. 4c). For broader lists like the GWAS and OMIM catalogs, we found no significant evidence of depletion or enrichment. We observed a similar pattern for multi-tissue eQTL genes (Extended Data Fig. 10b). Nonetheless, outlier expression affected some important and actionable disease genes. We observed multi-tissue outliers for five ACMG genes, five high-risk cardiovascular disease genes, and 14 cancer genes (Extended Data Table 2, Supplementary Table 1). However, the direction of the known disease-causing mutations and of the expression in the outlier individual were only consistent for a subset of the genes. Of the 20 unique genes from these lists, five had an underexpression outlier when the disease was caused by loss of the gene’s function. For example, one individual was an underexpression outlier (median Z-score of -3.5) for *SDHAF2*, a tumor suppressor on the ACMG list. *SDHAF2* promotes assembly of the SDH complex, which functions in the citric acid cycle and the electron transport chain. Loss of *SDHAF2* function impairs cellular respiration, which triggers hypoxia signalling and leads to the development of paragangliomas (neuroendocrine tumors)^27^,^28^. In addition to finding outliers in these highly curated disease gene lists, we found multi-tissue outliers for 784 ClinVar, 770 OrphaNet, 813 OMIM and 1532 GWAS genes compared to the expectation under independence of 734, 734, 762, and 1608, respectively (Extended Data Table 2, Supplementary Table 1). These results suggest that, although disease genes are depleted for multi-tissue outliers, due in part to selective pressure, extreme expression may contribute to disease risk.

### Expression data improves variant prioritization

In addition to characterizing the regulatory impact of rare variation across the GTEx cohort in aggregate, we sought to prioritize candidate regulatory variants from each individual genome. Existing methods for predicting rare variant impact use epigenetic data and other genomic annotations derived from external studies^9^–^13^. We hypothesized that by integrating gene expression data from the same individual whose genome we seek to analyze, along with these external annotations, we could significantly improve our identification of rare regulatory variants. We developed RIVER (RNA-Informed Variant Effect on Regulation), a probabilistic modeling framework that jointly analyzes personal genome and transcriptome data to estimate the probability that a variant has regulatory impact in that individual (https://github.com/ipw012/RIVER, see Online methods). RIVER is based on a generative model that assumes that genomic annotations (Extended Data Table 3), such as the location of a variant with respect to regulatory elements, determine the prior probability that variant is a *functional regulatory variant*, which is an unobserved variable. The functional regulatory variant status then influences whether nearby genes are likely to display outlier levels of gene expression in that person (Fig. 5a). RIVER is trained in an unsupervised manner. It does not require a labeled set of functional/non-functional variants; rather it derives its predictive power from identifying expression patterns that tend to coincide with particular rare variant annotations.

**Figure 5.**
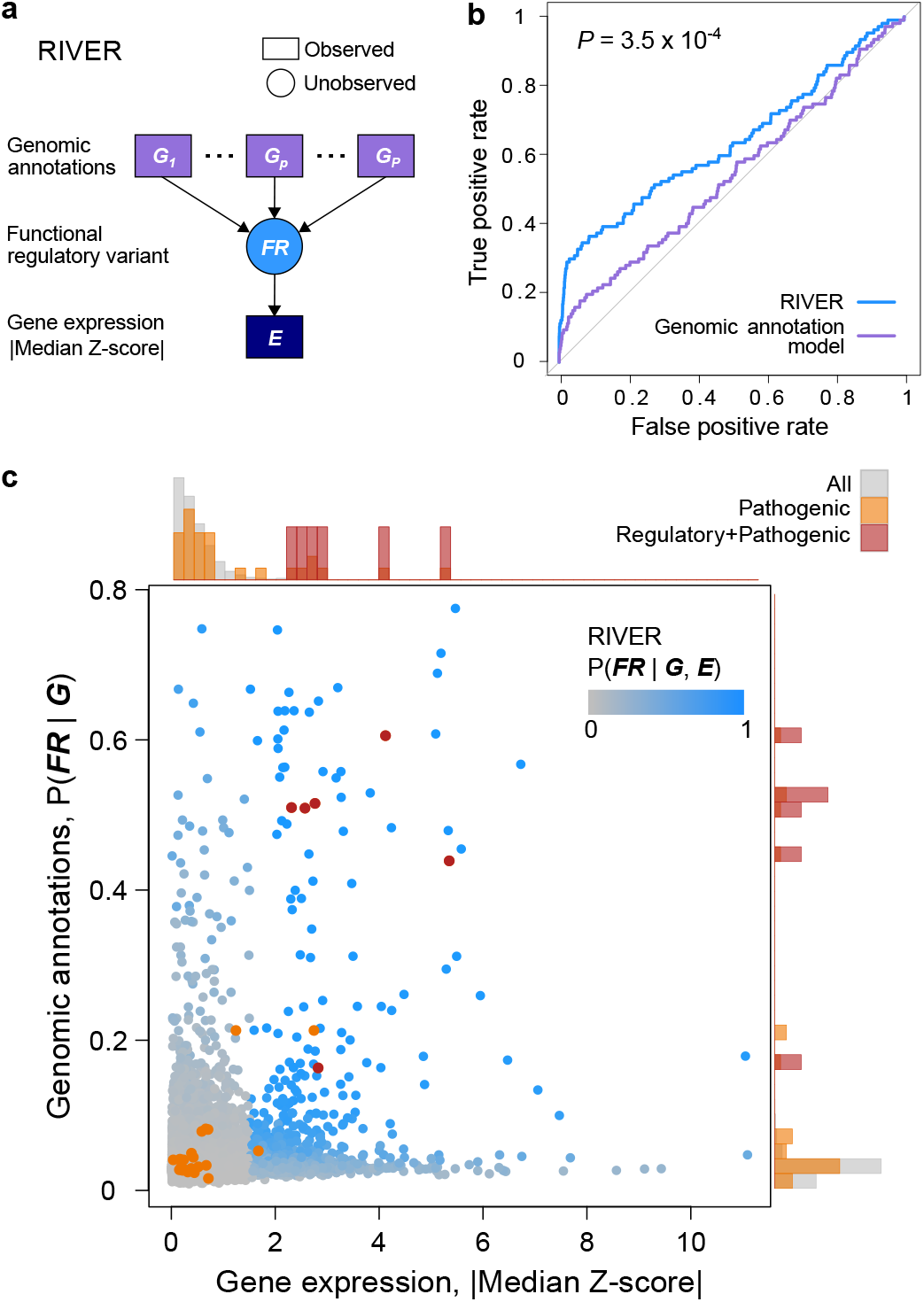
Performance of RIVER for prioritizing functional regulatory variants. (a) RIVER probabilistic graphical model (see Online methods). (b) Predictive power of RIVER compared with a L2-regularized logistic regression model using only genomic annotations without gene expression data. Accuracy was assessed using matched, held out individuals sharing the same rare variants as observed individuals (see Online methods, AUCs compared with DeLong’s approach^29^). (c) Distribution of RIVER scores (shades of blue) as a function of scores from genomic annotation or gene expression alone. Pathogenic SNVs annotated in ClinVar are shown in red if they were likely regulatory (nonsense, splice-site, or synonymous) and orange otherwise (missense). The distributions of variant categories across absolute median Z-scores and predictions from genomic annotation are shown as histograms aligned opposite the corresponding axes.

We applied RIVER to the GTEx cohort, training model parameters using 48,575 instances where an individual had at least one rare variant within 10 kb of the TSS of a gene with expression observed in at least five tissues. Here we assessed outliers using a more lenient threshold of |median Z-score| ≥ 1.5 with no restriction to a single individual per gene, both to obtain a larger set of outliers for training and to allow application to larger cohorts or external test instances. For evaluation, we held out pairs of individuals at genes where only those two individuals shared the same rare variants. We then computed the RIVER score (the posterior probability of having a functional regulatory variant given both whole genome and RNA sequencing data) from one individual, and assessed the accuracy with respect to the second individual’s held out expression levels (see Online methods). Using this labeled test data, we evaluated the predictive accuracy of RIVER compared with a L2-regularized multivariate logistic regression model that uses genomic annotations alone, and observed a significant improvement by incorporating expression data (Fig. 5b, AUC for RIVER and the genomic annotation model were 0.638 and 0.541, respectively, *P* = 3.5 × 10^−4^). Allele-specific expression was also enriched among the top RIVER instances compared with genome annotation models (Extended Data Fig. 11). Although RIVER was trained in an unsupervised manner, the learned model prioritized variants that were supported by both extreme expression levels for a nearby gene and genomic annotations suggestive of potential impact (Fig. 5c). Rather than using a heuristic or manual approach, RIVER automatically learns the relationship between genomic annotations and changes in gene expression from data to provide a coherent estimate of the probability of regulatory impact. For instance, multi-tissue outliers with a large proportion of single-tissue outliers were more likely to have high RIVER scores (Extended Data Fig. 12). Using a simplified supervised model, we estimated that even after accounting for the most informative genomic annotations or summary scores from state-of-the-art models including CADD and DANN, an individual was more likely to be an expression outlier if another individual with matched rare variants was an outlier (average log-odds ratio 2.76, Extended Data Table 4). This simplified approach supported the benefit of integrating gene expression data into variant prioritization.

To investigate how RIVER might inform disease variant analysis, we intersected rare variants in the GTEx individuals with variants from ClinVar^30^ (Extended Data Table 5). We identified 27 pathogenic or risk variants present in 21 individuals, and evaluated the RIVER score of each (Fig. 5c). Overall, pathogenic variants scored higher than background variants (two-sided Wilcoxon rank sum test, *P* = 3.25 × 10^−9^, Extended Data Fig. 13). We note that rare indels and SVs were not found nearby the genes in the individuals carrying these pathogenic variants. Considering that ClinVar is biased toward protein-coding variants, we observed that six of the 27 variants were annotated as nonsense, splice site, or synonymous variants, with the rest being missense. These likely regulatory variants had RIVER scores of 0.980 on average, putting them in the top 99.9^th^ percentile. Among these, three individuals harbored the minor allele at two distinct pathogenic variants (rs113993991 and rs113993993) near *SBDS*, each associated with Shwachman-Diamond syndrome. This recessive syndrome causes systemic symptoms including pancreatic, neurological, and hematologic abnormalities^31^ and can disrupt fibroblast function^32^. The GTEx individuals were heterozygous for these variants and thus lacked the disease phenotype. Nonetheless, we saw extreme underexpression of *SBDS* across almost all tissues in these individuals, including brain tissues, fibroblasts, and pancreas (Extended Data Fig. 14). In another case, an individual harbored the minor allele of rs80338735, which is associated with cerebral creatine deficiency syndrome 2, shown to cause neurological deficiencies and also lead to low body fat^33^. The nearby gene *GAMT* showed the most extreme underexpression (Z-score < -4) in adipose (subcutaneous), although unfortunately no brain tissue was available for evaluation in this individual (Extended Data Fig. 14). These cases demonstrate that RIVER can provide an important and novel ability to prioritize disease-causing regulatory variants by integrating population-scale patterns of gene expression.

## Discussion

Using whole genome sequences and RNA-seq from 44 human tissues from the GTEx project, we identified high-confidence gene expression outliers and completed the largest study to date of rare variants impacting gene expression. Outliers were better explained by genetic variation when we combined expression data from multiple tissues. We found that rare structural variants, frameshift indels, coding variants, and variants near the transcription start site were most likely to have large effects on expression. These effects were often directional; for example, we saw duplications tended to cause overexpression while deletions and stop-gain variants caused underexpression. Rare coding variants can alter expression as much as non-coding variation, underscoring that expression data can reveal the molecular consequences of disrupting protein-coding regions. Consistent with the effects of purifying selection, less constrained genes were most likely to have multi-tissue outliers. However, we discovered examples of outliers for known disease genes where the previously known causal variant was protein-coding and associated with altered expression. This highlights the benefit of population-scale functional genomics data for both non-coding and coding variant interpretation.

Building on observations from the population analysis, we developed RIVER, which combines individual RNA-seq and whole genome data to predict which rare variants have large regulatory impact. Integrating genomic features with expression dramatically improved variant prioritization, supporting that gene expression data can help interpret variants with effects that are unclear from genome sequence alone. RIVER is an extensible predictive model for combining whole genome sequences with molecular phenotypes to identify high-impact variants. Therefore, our results suggest a general approach that can be applied to studies supplementing genomes with other molecular phenotypes, such as methylation^34^–^36^ and histone modification^37^,^38^. We anticipate that such integrative approaches will be essential for effective interpretation of genome-wide genetic variation on a personalized level.

## Online methods

### Study population

All human subjects were deceased donors. Informed consent was obtained for all donors via next-of-kin consent to permit the collection and banking of de-identified tissue samples for scientific research.

### Whole genome sequence and multi-tissue RNA-seq data

Whole genome sequencing (WGS) VCF files from the GTEx V6 release were downloaded from the dbGaP. In addition, we downloaded RPKM, read counts, and allele-specific expression data comprising 8,555 RNA-seq samples from 53 tissues and 544 individuals. Of these individuals, 520 have whole exome sequencing and 148 have additional whole genome sequencing.

### Correction for technical confounders

We restricted our expression analyses to the 449 individuals and 44 tissues for which sex and the top three genotype principal components, which capture major population stratification, were available. For each tissue, we log_2_-transformed all expression values (RPKM + 2). We then standardized the expression of each gene to encourage normality while preventing shrinkage of outlier expression values caused by quantile normalization. To remove unmeasured batch effects and other confounders, for each tissue separately, we estimated hidden factors using PEER^21^ on the transformed expression values. In each tissue, we defined expressed genes as those with at least 10 individuals with RPKM > 0.1 and read count > 6. Genes falling below these thresholds were discarded from the analysis for that tissue. The number of PEER factors estimated per tissue was determined by sample size (N): 15 factors for N < 150, 30 factors for 150 ≤ N ≤ 250, and 35 factors for N > 250. We regressed out the PEER factors, the top three genotype principal components, and sex (where appropriate) from the transformed expression data for each tissue using linear regression. Finally, we standardized the expression residuals for each gene, which yielded Z-scores.

### Single-tissue and multi-tissue outlier discovery

Single-tissue and multi-tissue outlier calling was restricted to autosomal lincRNA and protein coding genes. In addition to the constraints described in the main text, we only tested for multi-tissue outliers among individuals with expression measurements for the gene in at least five tissues. To reduce cases where non-genetic factors may cause widespread outliers, we removed eight individuals that were multi-tissue outliers for 50 or more genes from all downstream analyses. These individuals were also removed before single-tissue outlier discovery.

### Replication of expression outliers

We evaluated the replication of single-tissue outliers between pairs of tissues. We calculated the proportion of outliers discovered in one tissue that had a |Z-score| ≥ 2 for the same gene in the replication tissue. We also required that the replication Z-score have the same sign as the Z-score in the discovery tissue. Since each tissue had a different number of samples and certain groups of tissues were sampled in a specific subset of individuals, we evaluated the extent to which replication was influenced by the size and the overlap of the discovery and replication sets. To make pairs of tissues comparable, we repeated the replication analysis with the discovery and replication in exactly 70 individuals for each pair of tissues with enough overlapping samples. We compared the replication patterns in this subsetted analysis to those obtained by using all individuals for discovery and replication. To estimate the extent to which individual overlap biased replication estimates, for each pair of tissues with a sufficient number of samples, we defined three disjoint group of individuals: 70 individuals with data for both tissues, 69 distinct individuals with data in the first tissue, and 69 distinct individuals with data in the second tissue. We discovered outliers in the first tissue using the shared set of individuals. Then we tested the replication of these outliers in the discovery individuals in the second tissue. Finally, for each gene, we added the identified outlier to distinct set of individuals and tested the replication again in the second tissue. We repeated the process running the discovery in the second tissue and the replication in the first one. We compared the replication rates when using the same or different individuals for the discovery and replication.

We assessed the confidence of our multi-tissue outliers using cross-validation. Specifically, we separated the tissue expression data randomly into two groups: a discovery set of 34 tissues and a replication set of 10 tissues. For *t* = 10, 15, 20, 25, and 30, we randomly sampled *t* tissues from the discovery set and performed outlier calling as described above. To assess the replication rate, we computed the proportion of outliers in the discovery set with |median Z-score| ≥ 1 or 2 in the replication set. We set no restriction on the number of tissues required for testing in the replication set. To calculate the expected replication rate, we randomly selected individuals in the discovery set requiring that each individual show expression in at least five tissues for the gene. We then computed the replication rate for this background using the procedure described above. We repeated this process 10 times for each discovery set size.

### Quality control of genotypes and rare variant definition

We restricted our rare variant analyses to individuals of European descent, as they constituted the largest homogenous population within our dataset. We considered only autosomal variants that passed all filters in the VCF (those marked as PASS in the Filter column). Minor allele frequencies (MAF) within the GTEx data were calculated from the 123 individuals of European ancestry with whole genome sequencing data. The MAF was the minimum of the reference and the alternate allele frequency where the allele frequencies of all alternate alleles were summed together. Rare variants were defined as having MAF ≤ 0.01 in GTEx, and for SNVs and indels we also required MAF ≤ 0.01 in the European population of the 1000 Genomes Project Phase 3 data^20^. We also sought to ensure that population structure among the individuals of European descent was unlikely to confound our results. Therefore, we verified that the allele frequency distribution of rare variants included in our analysis (within 10 kb of a protein coding or lincRNA gene, see below) was similar for the five European populations in the 1000 Genomes project (Extended Data Fig. 1b).

### Enrichment of rare and common variants near outlier genes

We assessed the enrichment of rare SNVs, indels, and SVs near outlier genes. Proximity was defined as within 10 kb of the TSS for all analyses, with the exception of Fig. 3 where we included all variants within 10 kb of the gene, including the gene body, (200 kb for enhancers and SVs) to also capture coding variants. For each gene with a multi-tissue outlier, we chose the remaining set of individuals tested for multi-tissue outliers at the same gene as our set of non-outlier controls. We only included genes that had both a multi-tissue outlier and at least one control. We stratified variants of each class into four minor allele frequency bins (0–1%, 1–5%, 5–10%, 10–25%) to compare the relative enrichments of rare and common variants. We also assessed the enrichment of SNVs at different Z-score cutoffs. Enrichment was defined as the ratio of the proportion of outliers with a rare variant within 10 kb of the transcription start site (TSS) to the proportion of non-outliers with a rare variant in the same window. This enrichment metric is equivalent to the relative risk of having a nearby rare variant given outlier status. We used the asymptotic distribution of the log relative risk to obtain 95% Wald confidence intervals. Within our set of European individuals, we observed some individuals with minor admixture that had relatively more rare variants than the rest (Extended Data Fig. 2b). We confirmed that inclusion of these admixed individuals did not substantially affect our results (Extended Data Fig. 2c). We also calculated rare variant enrichments when restricting to variants outside protein-coding and lincRNA exons in Gencode v19 annotation (extending internal exons by 5 bp to capture canonical splice regions).

To measure the informativeness of variant annotations (Extended Data Table 1), we used logistic regression to model outlier status as a function of the feature of interest, which yielded log odds ratios with 95% Wald confidence intervals. Note that for the feature enrichment analysis in Fig. 3b and Extended Data Fig. 9, we required that outliers and their gene-matched non-outlier controls have at least one rare variant near the gene. We scaled all features, including binary features, to have mean 0 and variance 1 to facilitate comparison between features of different scale. We also calculated the proportion of overexpression outliers, underexpression outliers and non-outliers with a rare variant near the gene TSS (within 10 kb for SNVs and indels and 200kb for SVs). To each outlier instance, we assigned at most one of the 12 rare variant classes we considered, which are listed in Fig. 3. If an outlier had rare variants from multiple classes near the relevant genes, we selected the class that was most significantly enriched among outliers.

### Annotation of variants

We obtained annotations for SV categories from Chiang et al.^22^. We computed features for rare SNVs and indels using three primary data sources: Epigenomics Roadmap^39^, CADD v1.2^9^, and VEP v80^40^. We downloaded the promoter and enhancer annotation tracks produced by the Epigenomics Roadmap Project (http://www.broadinstitute.org/~meuleman/reg2map/HoneyBadger2_release/). We mapped 28 unique tissues in the GTEx Project to 19 tissue groups in the Roadmap Project. Using these annotations, for each individual, we then assessed whether each SNV or indel overlapped a promoter or enhancer region in at least one of the 19 Roadmap tissue groups. We obtained features including conservation, chromatin segmentation, and deleteriousness from the full annotation tracks of the CADD v1.2 release (downloaded 15/05/2015; http://cadd.gs.washington.edu/download). We intersected the rare variants segregating in each individual with the CADD annotation and extracted the conservation features. Finally, we obtained protein-coding and transcription-related annotations from VEP. This information was provided in the GTEx V6 VCF file. Using the pipeline described above, we generated features at the site-level for all 123 European individuals with WGS data. We then collapsed these features to generate gene-level features. The collapsed features are described in Extended Data Tables 1 and 3.

### Allele-specific expression (ASE)

We only considered sites with at least 30 total reads and at least five reads supporting each of the reference and alternate alleles. To minimize the effect of mapping bias, we filtered out sites that showed mapping bias in simulations^41^, that were in low mappability regions (ftp://hgdownload.cse.ucsc.edu/gbdb/hg19/bbi/wgEncodeCrgMapabilityAlign50mer.bw), or that were rare variants or within 1 kb of a rare variant in the given individual (the variants were extracted from GTEx exome sequencing data). The first two filters were provided in the GTEx ASE data release (Aguet *et al.,* GTEx cis-eQTL paper, co-submitted). The third filter was applied to eliminate potential mapping artefacts that mimic genetic effects from rare variants. We measured ASE effect size at each testable site as the absolute deviation of the reference allele ratio from 0.5. For each gene, all testable sites in all tissues were included. We compared ASE in single-tissue and multi-tissue outliers at different Z-score thresholds to non-outliers using a two-sided Wilcoxon rank sum test. To obtain a matched background, we only included a gene in the comparison when ASE data existed for both the outlier individual and at least one non-outlier. In the case of single-tissue outliers, we also required the tissue to match between the outlier and the non-outlier. All individuals that were neither multi-tissue outliers for the given gene nor single-tissue outliers for the gene in the corresponding tissue were included as non-outliers.

### Allele frequency measurements in UK10K

UK10K^3^ VCF files of whole genome cohorts were downloaded from https://www.ebi.ac.uk. We merged the Avon Longitudinal Study of Parents and Children (ALSPAC) EGAS00001000090 and the Department of Twin Research and Genetic Epidemiology (TWINSUK) EGAS00001000108 datasets for a total of 3,781 individuals. We counted the occurrence of all rare GTEx SNVs in Epigenomics Roadmap-annotated promoter regions among the UK10K samples. GTEx variants absent from the UK10K cohorts were assigned a count of 0.

### Enrichment of genes with multi-tissue outliers as eGenes

We defined multi-tissue eGenes using two approaches. The first, referred to as the tissue-by-tissue approach, considered cis-eQTL effects individually for each tissue and ignored sharing of effects between tissues. For this first approach, we obtained lists of significant eGenes (FDR ≤ 0.05) for each of the 44 tissues from the GTEx V6p release. The second approach defined cis-eQTLs with shared effects across tissues using the RE2 model as part of the Metasoft software^42^. The GTEx Consortium performed shared cis-eQTL discovery using this model for 19,172 autosomal lincRNA and protein coding genes with cis-eQTLs passing the significance level (FDR ≤ 0.05) in at least one of the 44 tissues in the tissue-by-tissue analysis (Aguet *et al.*, GTEx cis-eQTL paper, co-submitted). Using these results, we chose for each gene the variant with the lowest nominal *P*-value from the RE2 model. We then determined the number of tissues in which this variant-gene pair showed a cis-eQTL effect; we used as our threshold an m-value ≥ 0.9^42^. For each of the 18,380 genes tested for multi-tissue outliers, we calculated the number of tissues in which the gene appeared as a significant eGene (approach 1) or had a shared eQTL effect (approach 2). We compared this value for genes with and without a multi-tissue outlier with a two-sided Wilcoxon rank sum test.

Finally, we wanted to show that this enrichment of outlier genes as multi-tissue eGenes was not confounded by gene expression level. To this end, using the Metasoft results, we stratified genes tested for multi-tissue outliers into RPKM deciles and repeated the comparison between genes with and without a multi-tissue outlier.

### Evolutionary constraint of genes with multi-tissue outliers

We obtained gene level estimates of evolutionary constraint from the Exome Aggregation Consortium^26^ (http://exac.broadinstitute.org/, ExAC release 0.3). By jointly analyzing the patterns of exonic variation in over 60,000 exomes, the ExAC database can be used to rank evolutionary constraint of genes based on their tolerance for synonymous, missense, and loss-of-function variation. We intersected the 17,351 autosomal lincRNA and protein coding genes with constraint data from ExAC with the 18,380 genes tested for multi-tissue outliers from GTEx, yielding 14,379 genes for further analysis (3,897 and 10,482 genes with and without an outlier, respectively). We examined three functional constraint scores from the ExAC database: synonymous Z-score, missense Z-score, and probability of loss-of-function intolerance (pLI). We defined sets of synonymous and missense intolerant genes as genes with a synonymous or missense Z-score above the 90^th^ percentile. We defined loss-of-function intolerant genes as those with a pLI score above 0.9 following the guidelines provided by the ExAC consortium. We then tested for the enrichment of genes with multi-tissue outliers in the lists of synonymous, missense, and loss-of-function intolerant genes. We calculated odds ratios and 95% confidence intervals using Fisher’s exact test. We repeated this analysis for three other gene lists: 19,172 multi-tissue eGenes from GTEx V6p defined using Metasoft^42^, 9,480 reported GWAS genes from the NHGRI-EBI catalog^43^ (http://www.ebi.ac.uk/gwas accessed 30/11/2015), and 3,576 OMIM genes (http://omim.org/ accessed 26/5/2016). Multi-tissue eGenes were ranked by nominal *P*-value from the RE2 random effects model implemented in Metasoft. The top 3,879 multi-tissue eGenes were classified as shared eGenes, while the remaining 11,386 genes were considered as a background. The number of shared eGenes was chosen to match the number of multi-tissue outlier genes in the intersection with the ExAC database.

We tested for a difference in the mean constraint for genes with multi-tissue outliers and genes with multi-tissue eQTLs using ANOVA. For each of the three constraint scores in ExAC, we treated the score for each gene as the response and the status of the gene as having a multi-tissue outlier and/or a multi-tissue eQTL as a categorical predictor with four classes. After fitting the model, we performed Tukey’s range test to determine whether there was a significant difference in the mean constraint between genes with a multi-tissue outlier but no multi-tissue eQTL and genes with a multi-tissue eQTL but no multi-tissue outlier.

### Overlap of genes with multi-tissue outliers and disease genes

We examined the enrichment of genes with multi-tissue outliers in eight disease gene lists. A summary of these lists can be found in Extended Data Table 2. We tested for enrichment or depletion by comparing each disease gene list to the 18,380 genes tested for multi-tissue outliers. We computed odds ratios and 95% confidence intervals using Fisher’s exact test. We compared the enrichments for genes with multi-tissue outliers to genes with shared eQTL effects as defined by the Metasoft approach. Like the ExAC enrichment analysis, we chose the top 4,919 shared eGenes (ranked by *P*-value from the RE2 model) to match the number of genes with multi-tissue outliers.

Heritable cancer predisposition and heritable cardiovascular disease gene lists were curated by local experts in clinical and laboratory-based genetics in the two respective areas (Stanford Medicine Clinical Genomics Service, Stanford Cancer Center's Cancer Genetics Clinic, and Stanford Center for Inherited Cardiovascular Disease). Genes were included if both the clinical and laboratory-based teams agreed there was sufficient published evidence to support using variants in these genes in clinical decision making.

### RIVER integrative model for predicting regulatory effects of rare variants

RIVER (RNA-Informed Variant Effect on Regulation) is a hierarchical Bayesian model that predicts the regulatory effects of rare variants by integrating gene expression with genomic annotations. The RIVER model consists of three layers: a set of nodes *G = G_1_ … G_P_* in the topmost layer representing *P* observed genomic annotations over all rare variants near a particular gene, a latent binary variable *FR* in the middle layer representing the unobserved functional regulatory status of the rare variants, and one binary node *E* in the final layer representing expression outlier status of the nearby gene. We model each conditional probability distribution as follows:

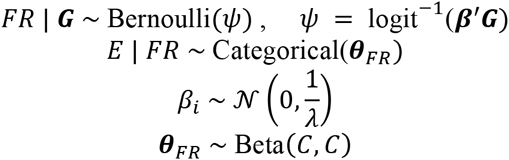

with parameters *β* and *θ* and hyper-parameters *λ* and *C.*

Because *FR* is unobserved, the RIVER log-likelihood objective over instances *n = 1, …, N* 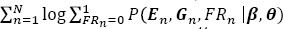 is non-convex. We therefore optimize model parameters via Expectation-Maximization^44^ (EM) as follows:

In the E-step, we compute the posterior probabilities 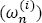 of the latent variables *FR_n_* given current parameters and observed data. For example, at the *i*th iteration, the posterior probability *of FR_n_ =* 1 for the *n*th instance is

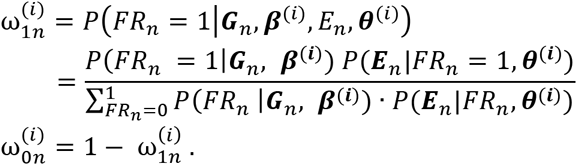

In the M-step, at the *i*th iteration, given the current estimates ω^(i)^, the parameters (*β*^*(i + 1)*^*) are estimated as

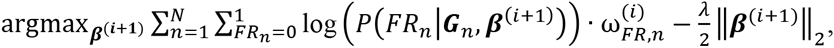

where *λ* is an L2 penalty hyper-parameter derived from the Gaussian prior on *β*.

The parameters *θ* get updated as:

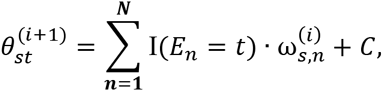

where I is an indicator operator, *t* is the binary value of expression *E_n_, s* is the possible binary values of *FR_n_*, and *C* is a pseudo count derived from the Beta prior on *θ*. The E and M steps are applied iteratively until convergence.

### RIVER application to the GTEx cohort

As input, RIVER requires simply a set of genomic features ***G*** and a set of corresponding expression outlier observations ***E***, each over a set of instances representing one gene in one individual. Using the variant annotations described above, we generated genomic features at the site-level for the 116 European individuals with GTEx WGS data that had fewer than 50 multi-tissue outliers. We then collapsed these features for all rare SNVs within 10 kb of each TSS to generate gene-level features with relevant computational operators: a binary indicator implying a presence/absence of any rare SNV in each of the VEP features, total number of rare SNVs in each of the VEP features, chromatin states from ChromHMM, and Segway segmentations, and a maximum value over all nearby rare SNVs for the rest of the features. The collapsed features are described in Extended Data Table 3. This produces a matrix of genomic features ***G*** of size (116 individuals × 1,736 genes) × (112 genomic features), which we standardize within features (columns) before use. The corresponding multi-tissue outlier values ***E*** were computed using PEER-corrected gene expression Z-scores from each tissue. For each gene, we defined any individual with |median Z-score| ≥ 1.5 as an outlier if the expression was observed in at least five tissues; the remaining individuals were labeled as non-outliers for the gene. In total, we extracted 48,575 instances where an individual had at least one rare variant within 10 kb of TSS of a gene. We then incorporated standardized genomic features (***G*** nodes in Fig. 5a) and multi-tissue outlier states (***E*** node in Fig. 5a) as input to RIVER.

To train and evaluate RIVER on the GTEx cohort, we first identified 3,766 instances of individual and gene pairs where two individuals had the same rare SNVs near a particular gene. We used these instances for evaluation as described below. We held out those instances and trained RIVER parameters with the remaining instances. RIVER requires two hyper-parameters *λ* and *C*. To select *λ*, we first applied a multivariate logistic regression with features ***G*** and response variable ***E***, selecting lambda with the minimum squared error via 10-fold cross-validation (we selected *λ* = 0.01). We selected *C* = 50, informed simply by the total number of training instances available, as validation data was not available for extensive cross-validation. Initial parameters for EM were set to ***θ*** = (P(*E* = 0 | *FR* = 0), P(*E* = 1 | *FR* = 0), P(*E* = 0 | *FR* = 1), P(*E* = 1 | *FR* = 1)) = (0.99, 0.01, 0.3, 0.7) and ***β*** from the multivariate logistic regression above, although different initializations did not significantly change the final parameters (Extended Data Table 6).

The 3,766 held out pairs of instances from individuals with an identical rare variant were used to create a labeled evaluation set. For one of the two individuals from each pair, we estimated the posterior probability of a functional rare variant P(*FR* | ***G***, ***E***, ***β***, ***θ***). The outlier status of the second individual, whose data was not observed either during training or prediction, was then treated as a “label” of the true status of functional effect *FR*. Using this labeled set, we compared the RIVER score to the posterior P(*FR* | ***G, β***) estimated from the plain multivariate logistic regression model with genomic annotations alone. We produced ROC curves and computed AUC for both models, testing for significant differences using DeLong’s method^29^. This metric relies on outlier status reflecting the consequences of rare variants— pairs of individuals who share rare variants tend to have highly similar outlier status even after regressing out effects of common variants (Kendall’s tau rank correlation, *P* < 2.2 × 10^−16^). As a second metric, we also evaluated performance of both the genomic annotation model and RIVER by assessing ASE. We tested the association between ASE and model predictions using Fisher's Exact Test. High allelic imbalance, defined by a top 10% threshold on median absolute deviation of the reference-to-total allele ratio from an expected ratio (0.5) across 44 tissues, was compared to posterior probabilities of rare variants being functional from both models with four different thresholds (top 10% – 40%).

### Supervised model integrating expression and genomic annotation

To assess the information gained by incorporating gene expression data in the prediction of functional rare variants, we applied a simplified supervised approach to a limited dataset. We used the instances where two individuals had same rare variants to create a labeled training set where the outlier status of the second individual was used as the response variable. We then trained a logistic regression model with just two features: 1) the outlier status of the first individual and 2) a single genomic feature value such as CADD or DANN. We estimated parameters from the entire set of rare-variant-matched pairs using logistic regression to determine the log odds ratio and corresponding *P*-value of expression status as a predictor. While this approach was not amenable to training a full predictive model over all genomic annotations jointly, given the limited number of instances, it provided a consistent estimate of the log odds ratio of outlier status. We tested five genomic predictors: CADD, DANN, transcription factor binding site annotations, PhyloP scores, and one aggregated feature, posterior probability from a multivariate logistic regression model learned with all genomic annotations (Logistic) (Extended Data Table 4).

### RIVER assessment of pathogenic ClinVar variants

We downloaded pathogenic variants from the ClinVar database^30^ (accessed 04/05/2015). We searched for the presence of any of these disease variants within the set of rare variants segregating in the GTEx cohort. Using the ClinVar database, we then manually curated this set of variants, classifying them as pathogenic only if there was supporting clinical evidence of their role in disease. Specifically, any disease variant reported as pathogenic, likely pathogenic, or a risk factor for disease was considered pathogenic. To explore RIVER scores for those pathogenic variants, all instances were used for training RIVER. We then computed a posterior probability P(*FR* | ***G***, ***E***, ***β***, ***θ***) for each instance coinciding with a pathogenic ClinVar variant.

### Stability of estimated parameters with different parameter initializations

We tried several different initialization parameters for either ***β*** or ***θ*** to explore how this affected the estimated parameters. We initialized a noisy ***β*** by adding *K*% Gaussian noise compared to the mean of ***β*** with fixed ***θ*** (for *K* = 10, 20, 50 100, 200, 400, 800). For ***θ,*** we fixed P(*E* = 1 | *FR* = 0) and P(*E* = 0 | *FR* = 0) as 0.01 and 0.99, respectively, and initialized (P(*E* = 1 | *FR* = 1), P(*E* = 0 | *FR* = 1)) as (0.1, 0.9), (0.4, 0.6), and (0.45, 0.55) instead of (0.3, 0.7) with ***β*** fixed. For each parameter initialization, we computed Spearman rank correlations between parameters from RIVER using the original initialization and the alternative initializations. We also investigated how many instances within top 10% of posterior probabilities from RIVER under the original settings were replicated in the top 10% of posterior probabilities under the alternative initializations (Accuracy in Extended Data Table 6).

### Code availability

RIVER is available at https://github.com/ipw012/RIVER. Additionally, the code for running analyses and producing the figures throughout this manuscript is available separately (https://github.com/joed3/GTExV6PRareVariation).

## Data availability

The GTEx V6 release genotype and allele-specific expression data are available from dbGaP (study accession phs000424.v6.p1; http://www.ncbi.nlm.nih.gov/projects/gap/cgi-bin/study.cgi?study_id=phs000424.v6.p1). Expression data from the V6p release and eQTL results are available from the GTEx portal (http://gtexportal.org).

## Acknowledgements

We thank the MacArthur lab and the Laboratory, Data Analysis, and Coordinating Center (LDACC) for performing the quality control of the whole genome sequencing data. We also thank Donald Conrad for his help in generating the structural variant calls. The Genotype-Tissue Expression (GTEx) project was supported by the Common Fund of the Office of the Director of the National Institutes of Health (http://commonfund.nih.gov/GTEx). Additional funds were provided by the National Cancer Institute (NCI), National Human Genome Research Institute (NHGRI), National Heart, Lung, and Blood Institute (NHLBI), National Institute on Drug Abuse (NIDA), National Institute of Mental Health (NIMH), and National Institute of Neurological Disorders and Stroke (NINDS). Donors were enrolled at Biospecimen Source Sites funded by NCI\SAIC-Frederick, Inc. (SAIC-F) subcontracts to the National Disease Research Interchange (10XS170) and Roswell Park Cancer Institute (10XS171). The LDACC was funded through a contract (HHSN268201000029C) to The Broad Institute, Inc. Biorepository operations were funded through an SAIC-F subcontract to Van Andel Institute (10ST1035). Additional data repository and project management were provided by SAIC-F (HHSN261200800001E). The Brain Bank was supported by a supplement to University of Miami grant DA006227. E.K.T is supported by a Hewlett-Packard Stanford Graduate Fellowship and a doctoral scholarship from the Natural Science and Engineering Council of Canada. J.R.D. is supported by a Lucille P. Markey Biomedical Research Stanford Graduate Fellowship. J.R.D. and Z.Z. acknowledge the Stanford Genome Training Program (SGTP; NIH/NHGRI T32HG000044). Z.Z. is also supported by the National Science Foundation (NSF) GRFP (DGE-114747). F.N.D. is supported by the Joseph C. Pistritto Research Fellowship. B.J.S is supported by NIH training grant T32 GM007057. A.J.S. is supported by a Mr. and Mrs. Spencer T. Olin Fellowship for Women in Graduate Study. A.B. is supported by the Searle Scholars Program and NIH grant 1R01MH109905-01. A.B. and S.B.M. are supported by NIH grants R01MH101814 (NIH Common Fund; GTEx Program) and R01HG008150 (National Human Genome Research Institute (NHGRI); Non-Coding Variants Program). S.B.M. is supported by NHGRI grants U01HG007436 and U01HG009080. We thank the artists who created and made freely available the individual (http://cliparts.co/clipart/2353518) and organ graphics (http://www.allvectors.com/human-organs/) that we modified slightly to use in Fig. 1a. We thank David A. Knowles for code review.

## Author contributions

X.L., Y.K., E.K.T., J.R.D., A.B., and S.B.M. designed the study, performed analyses, and wrote the manuscript. Y.K., F.N.D., and A.B. developed RIVER. C.C., A.J.S, and I.M.H. provided the set of SVs. J.M. provided the lists of curated cancer and cardiovascular disease genes. Z.Z., B.J.S., and A.G. contributed analysis and feedback. All authors provided comments during study design, analyses, and writing.

